# Postnatal gene restoration in succinic semialdehyde dehydrogenase deficiency (SSADHD) reveals phenotype reversibility

**DOI:** 10.64898/2026.03.24.713250

**Authors:** Henry H.C. Lee, Gabrielle McGinty, Amanda Liebhardt, Zihe Zhang, Björn Welzel, Sheryl Anne D. Vermudez, Erland Arning, Rui Lin, Minh Nguyen, Didem Cakici, Timothy Yu, Clifford J. Woolf, Phillip L. Pearl, Guangping Gao, Mustafa Sahin, Alexander Rotenberg

## Abstract

Succinic semialdehyde dehydrogenase deficiency (SSADHD) is a rare autosomal recessive metabolic disorder due to loss-of-function *ALDH5A1* mutations impairing the catabolism of γ-aminobutyric acid (GABA), the major inhibitory neurotransmitter in the brain. In SSADHD, pathologic accumulation of GABA and its metabolic by-product γ-hydroxybutyrate (GHB) corresponds to a clinical syndrome dominated by developmental delay and epilepsy in half of patients with risk of sudden death in adolescence and adulthood. Brain-wide *ALDH5A1* gene replacement for SSADHD is unavailable, and whether such treatment will reverse the SSADHD phenotype is unknown. We developed an inducible mouse SSADHD model, *Aldh5a1^lox-STOP^*, enabling Cre-dependent *Aldh5a1* restoration to evaluate gene therapy feasibility. In the absence of SSADH, *Aldh5a1^lox-STOP^* mice exhibit hyperactivity and excessive serum GHB levels, culminating in death by ∼postnatal day 22, recapitulating the severe SSADHD condition. Systemic delivery of a blood-brain barrier (BBB)-penetrating adeno-associated virus (AAV) carrying a *Cre* gene to *Aldh5a1^lox-STOP^* mice leads to brain-wide SSADH restoration, serum GHB level reduction, normalization of hyperactivity, and substantial increase in survival. As a step toward clinical translation, we further assessed an AAV encompassing a functional native promoter (FLnP) of *ALDH5A1* tethered to its human coding sequence, namely AAV-FLnP-hALDH5A1. *Aldh5a1^lox-STOP^* mice were effectively rescued when treated with AAV-FLnP-hALDH5A1 packaged in the blood-brain barrier (BBB)-penetrating capsid PHP.eB. These findings provide preclinical proof that SSADH gene replacement therapy is feasible and potentially effective.

## INTRODUCTION

Succinic semialdehyde dehydrogenase deficiency (SSADHD) is an autosomal recessive disorder due to loss-of-function mutations of the *ALDH5A1* gene ^1–4^. SSADH is critical for catabolism of γ-aminobutyric acid (GABA), the major inhibitory neurotransmitter in the brain ^5^. In SSADHD, a pathologic accumulation of GABA and its metabolite γ-hydroxybutyrate (GHB) leads to a profound developmental impairment with autistic features, impulsive behaviors, ataxia, and disordered sleep ^6–9^. Approximately 10% of patients with SSADHD are nonverbal and wheelchair bound with spasticity. Paradoxically, despite elevated GABA concentrations, about 50% develop epilepsy, with sudden unexplained death in epilepsy (SUDEP) or death from uncontrolled seizures, potentially due to compensatory downregulation of GABA receptors.^10–14^.

Disease-modifying SSADHD treatments are an unmet need. Behavioral symptoms and epilepsy in SSADHD are drug-resistant ^15–17^. Three targeted drug trials in SSADHD have either resulted in intolerable toxicity or lack of clinical benefit ^15,18–20^. Experimental enzyme replacement therapy (ERT) has been attempted in mice, but therapeutic efficacy was limited due to several challenges including short recombinant protein half-life and limited accessibility to the brain ^21^. Gene replacement therapy, the focus of the present report, offers advantages over ERT in terms of brain-targeted restoration, potentially enabling a disease-modifying SSADHD treatment; however, empirical data to demonstrate the feasibility of such therapy are absent^10,22,23^.

A necessary step toward gene replacement therapy for SSADHD is its de-risking in an animal model, by first determining whether key features of the SSADHD phenotype are reversible. A second practical concern for potential SSADHD gene therapies is whether such treatment can be delivered by existing viral vectors. Adeno-associated virus (AAV), the most widely used vector for human gene therapy, exhibits suitable neuron and astrocyte tropism; however, it is unknown whether AAV-mediated gene replacement in brain is sufficient for SSADHD phenotype rescue^10,24^.

To enable proof-of-concept tests of gene replacement therapy for SSADHD, we constructed a novel inducible SSADHD mouse model: *Aldh5a1^lox-STOP^*. In this mouse, the endogenous *Aldh5a1* gene is intrinsically inactivated by an inserted cassette in the first intron with STOP codons in all reading frames, leading to early transcriptional termination that can be relieved “on-demand” by *Cre*-mediated recombination ^25^, to restore *Aldh5a1* activity to its endogenous level. This novel inducible *Aldh5a1* mouse model enables systematic tests of whether the SSADHD phenotype can be reversed by postnatal gene restoration. In parallel, as a step toward clinical translation, we developed a novel AAV vector aimed at driving transgene expression that mimics endogenous *ALDH5A1* expression to minimize off-target effects and also to minimize the risk of overexpression which may result in excess synaptic GABA clearance ^26^. That is, we designed a novel AAV construct encompassing a human native promoter of *ALDH5A1* tethered to a full-length functional coding sequence of human *ALDH5A1*.

## RESULTS

### Construction and characterization of *Aldh5a1^lox-STOP^* mice

We used an Easi-CRISPR/Cas9, single-strand DNA (ssDNA)-assisted approach ^27^ to insert a lox-STOP-lox cassette in the first intron of the *Aldh5a1* gene located at chromosome 13 of the mouse genome, creating an *Aldh5a1^lox-STOP^* mouse. To ensure obligatory splicing after cassette insertion, we engineered the splice acceptor site sequence motif directly preceding exon two into the lox-STOP-lox cassette in this ssDNA (Fig. 1a, Fig. S1a. Resultant pups were screened by genomic PCR, identifying one positive founder (Fig. 1b, S1b-c). Next-generation sequencing confirmed obligatory splicing and sequence integrity (Fig. S1d-e, S2a-e).

**Fig. 1.**
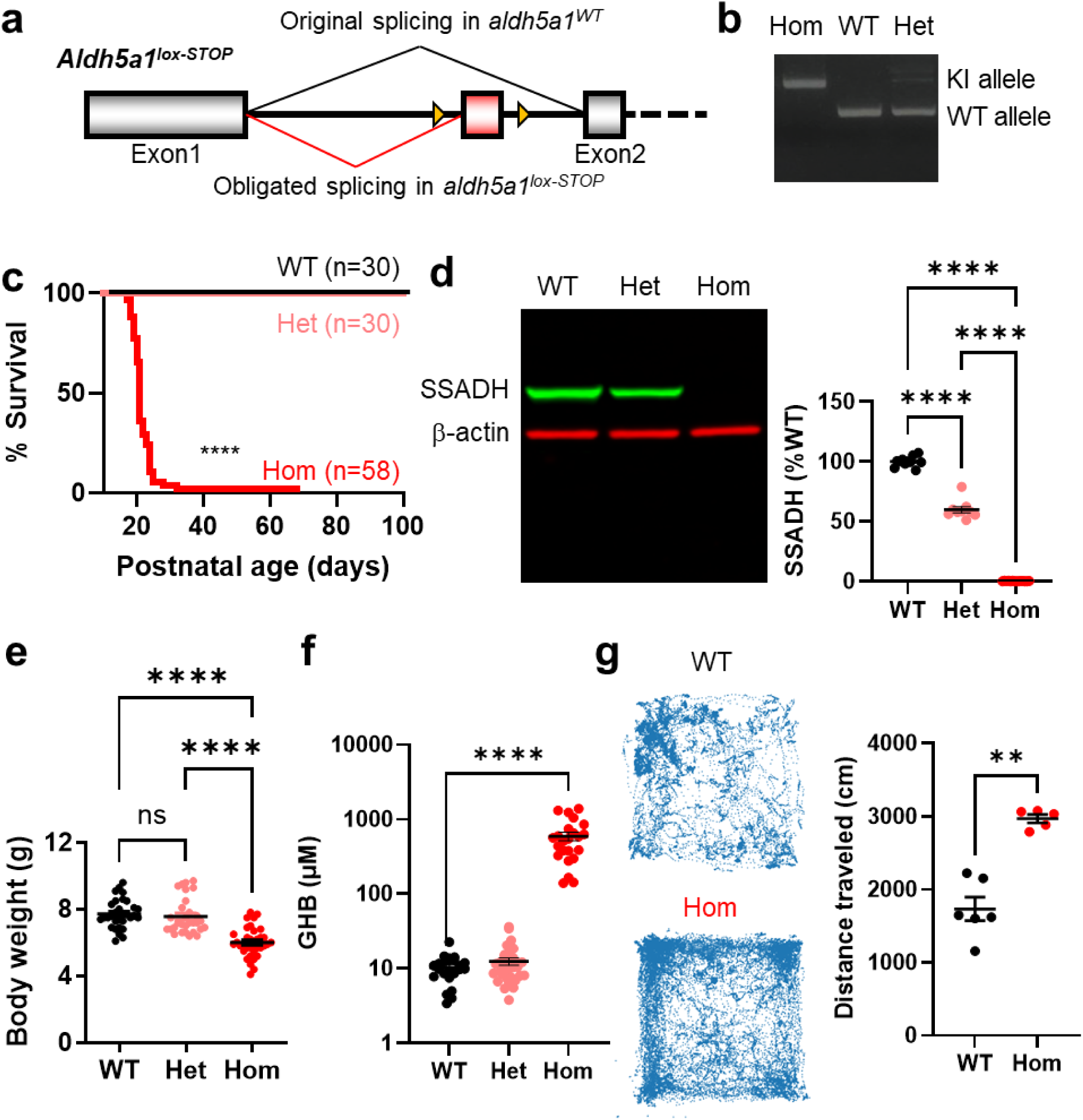
Construction and baseline characterization of the *Aldh5a1^lox-STOP^* mouse model. (a) Schematic diagram showing the insertion of a lox-STOP cassette in the first intron of the *Aldh5a1* gene, resulting in an obligated splicing and early termination of the translation of the endogenous *Aldh5a1* gene. (b) Genotype results showing the presence of knock-in (KI) and wild-type (WT) alleles in homozygous (Hom), heterozygous (Het), and WT littermates. (c) Survival curves of *Aldh5a1^lox-STOP^*WT, Het, and Hom up to 100 days of postnatal age. (d) Representative Western blot results showing the relative expressions of SSADH proteins in cortical lysates from WT, Het, and Hom mice at P16. β-actin is used as a loading control and for normalization. All data quantification is shown. (e) Body weight of WT, Het, and Hom mice at P16. Individual mouse data points are shown. (f) Serum GHB levels in WT, Het, and Hom mice at P16. (g) Free dwelling and exploratory behavior of WT and Hom mice. Value = MEAN ± SEM. **p<0.01, ****p<0.0001, ns=not significant, log-rank test in (c), One-way ANOVA followed by post-hoc Tukey’s multiple group comparisons in (d), (e), and (f). Mann-Whitney test in (g).

Obligated splicing in *Aldh5a1^lox-STOP^* should lead to total *Aldh5a1* gene inactivation in homozygous (Hom) *Aldh5a1^lox-STOP/lox-STOP^* mice. Accordingly, we confirmed that Hom mice have a severely restricted lifespan to postnatal day (P) 22.2 ± 0.9 in contrast to heterozygous (Het) *Aldh5a1^lox-STOP/WT^* mutants or wild-type (WT) *Aldh5a1^WT/WT^* control that were healthy and viable to at least P100 (n=30 WT, 30 Het, 58 Hom), without obvious signs of illness (Fig. 1c). Our Hom mice thus phenocopy a previously published *Aldh5a1* global knock-out (KO) mouse model ^28^ but enable *Cre*-dependent rescue (see results below). At a molecular level, we extracted total protein content from the cortical tissues of WT, Het, and Hom mice at P16. We found that Het mice have 59.88 ± 2.60 % SSADH expression compared to WT (p<0.0001, One-way ANOVA). In contrast, negligible amount of SSADH protein (0.23 ± 0.05 % WT, p<0.0001, One-way ANOVA) was detectable in all Hom mice (Fig. 1d, S3a). The body weight of WT, Het, and Hom mice at P16 were 7.71 ± 0.16 g, 7.57 ± 0.20 g, and 6.01 ± 0.15 g respectively (p<0.0001, One-way ANOVA) (Fig. 1e). Hom mice were universally underweight, regardless of sex (Fig. S4). We measured blood GHB content and found that WT and Het mice had serum concentrations of 10.29 ± 0.88 µM and 12.44 ± 1.36 µM GHB respectively (not significant, n.s.), but age-matched (P16) Hom mice have 592.20 ± 75.62 µM GHB, on average a >50-fold higher level relative to that in WT control (p<0.0001, One-way ANOVA) (Fig. 1f). We measured native exploratory and dwelling behaviors of *Aldh5a1^lox-STOP^* mice via an observer-independent, quantitative video approach described recently ^29^. High-speed video recording was used to capture native behaviors from the ventral position of a mouse^30^. The position and behaviors of mice were tracked inside the recording arena for 20 minutes. We found that Hom mice traveled 2969 ± 57 cm compared to 1734 ± 161 cm in WT mice (p=0.0043, Mann-Whitney) (Fig. 1g). Collectively, these data show that Hom mice at baseline are pathologic and developmentally delayed, with disease-relevant metabolic, molecular, behavior signatures, and high mortality in early life, recapitulating a severe SSADHD variant phenotype in humans.

### AAV-Cre-mediated brain-wide *Aldh5a1* restoration

Using a TdTomato reporter line, we confirmed *Cre*-dependent recombination broadly in the cortex and the hippocampus upon intraperitoneal (IP) injection of AAV-PHP.eB-Cre at a high dose (4 x 10^11^ gc/pup) ^31^ (Fig. 2a, b). To measure the impact of phenotype reversal upon *Aldh5a1* restoration, we injected AAV-PHP.eB-Cre into 29 P16 Hom mice and found that 18 survived beyond P30. Of those, 89% (16 out of 18) survived to at least P90 (p<0.0001, log-rank test) (Fig. 2c). Western blot analysis revealed that AAV-PHP.eB-Cre-rescued Hom mice had 68.57 ± 3.72% (Mean±SEM) SSADH protein content in their cortex compared to WT control at P90 at which time mice were euthanized for tissue harvest, a level comparable to untreated age-matched Het mice (60.41 ± 6.49% SSADH of WT, Mean±SEM, Fig. 2d). In the cortex of injected Hom mice that did not survive past P30, however, there was only 1.08 ± 0.42% SSADH protein (Mean±SEM, p<0.0001, One-way ANOVA). We also confirmed that SSADH proteins were stable in brain tissues collected post-mortem, validating the above results (Fig. S3b). The body weight of rescued Hom mice was 23.39 ± 0.91 g, compared to 21.85 ± 0.90 g in AAV-injected age-matched WT mice (Mean±SEM, ns, t-test) (Fig. 2e). Blood GHB content in AAV-PHP.eB-Cre-rescued Hom mice were 15.30 ± 1.51 µM compared to 10.29 ± 0.93 µM in AAV-injected WT control mice at P90 (Mean±SEM, p=0.0092, t-test, Fig. 2f). Upon brain-wide *Aldh5a1* restoration, AAV-PHP.eB-Cre-injected Hom mice traveled 1621 ± 208 cm compared to 1675 ± 133 cm as in WT control (Mean±SEM, ns, t-test, Fig. 2g). These results demonstrate that SSADHD phenotype is reversible upon brain-wide *Aldh5a1* restoration in Hom mice even if treatment commences at a symptomatic stage.

**Fig. 2.**
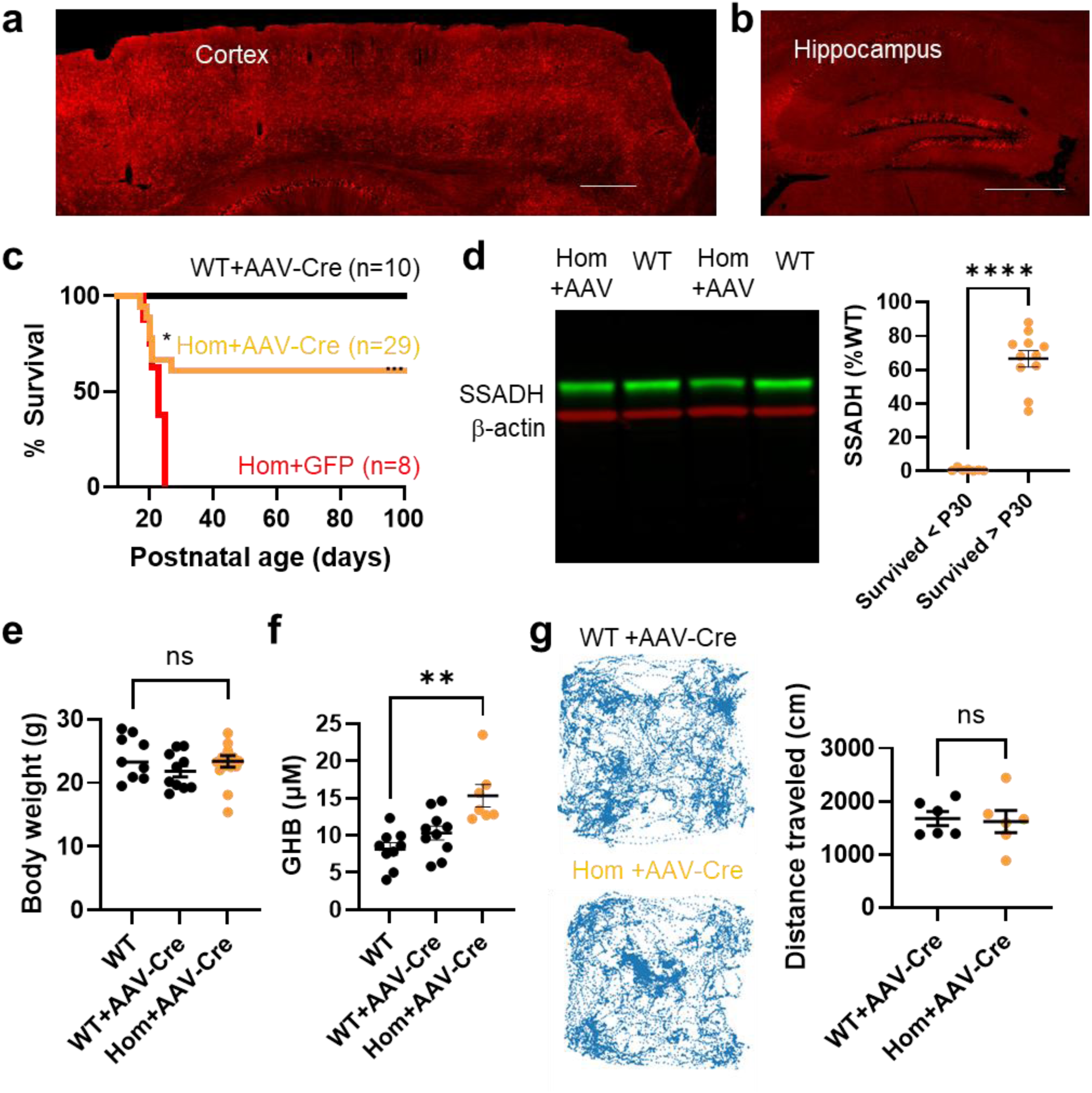
AAV-mediated brain-wide *Aldh5a1* restoration in *Aldh5a1^lox-STOP^* mice. Brain-wide transgene expression mediated by AAV-PHP.eB-CMV-Cre. (a-b) Confocal microscopy images of the cortex (a) and the hippocampus (b) sections from a Td-TOMATO mouse injected (via intraperitoneal injection) with 4×10^11^ gc AAV-CMV-Cre packaged in PHP.eB capsid. Scale bars: 0.4 mm. (c) Survival curve of wild-type (WT) or homozygous (Hom) *Aldh5a1^lox-STOP^* mice injected with AAV-Cre or AAV-GFP control vector, injected at P16. (d) Representative Western blot results of WT and Hom *Aldh5a1^lox-STOP^* mice upon AAV-Cre rescue. In those Hom+AAV mice but survived <P30, no SSADH expressions were found in their cortex. The amount of SSADH expressions (presented as %WT level) in those AAV-rescued Hom mice were compared with WT. (e) The body weight of AAV-Cre-injected WT and Hom at P100 are compared. Individual mouse data points are shown. (f) Serum GHB levels in WT and Hom treated with AAV-Cre, measured at P100. (g) Free dwelling and exploratory behavior of WT and Hom mice treated with AAV-Cre, measured at P100. Value = MEAN ± SEM. **p<0.01, ****p<0.0001, ns=not significant, log-rank test in (c), Mann-Whitney test in (d) and (g). One-way ANOVA followed by post-hoc multiple group comparisons in (e) and (f).

### AAV gene therapy candidate rescues *Aldh5a1^lox-STOP^* mice

We incorporated a native human *ALDH5A1* functional native promoter (FLnP, 1556 bp upstream sequence of transcriptional start) tethered with a human *ALDH5A1* coding sequence (1608 bp) into a PHP.eB capsid (namely AAV-PHP.eB-FLnP-ALDH5A1, or AAV-FLnP in short), and injected this at the same high dose (4 x 10^11^ gc/pup) in P16 mice (Fig. 3a). We injected 22 Hom mice (at P16) and found that 15 of them survived beyond P30. Of those, 60% (9 out of 15) survived beyond P90 (p<0.0001, log-rank test) (Fig. 3b). Western blot analysis revealed that AAV-FLnP-rescued Hom mice that survived between >P30 and <P90 (at which time mice were euthanized for tissue harvest) had 41.10 ± 11.17% SSADH protein content in their cortex compared to 154.07 ± 15.66 % WT SSADH protein in those that survived >P90, indicating a positive correlation between restored SSADH protein levels and survival in treated mice (R^2^=0.6825, p<0.0001) (Fig. 2c).The body weight of rescued Hom mice was 27.11 ± 1.51 g, compared to 27.47 ± 1.31 g in AAV-injected age-matched WT mice (ns, t-test) (Fig. 3c). Blood GHB content in AAV-PHP.eB-Cre-rescued Hom mice were 19.13 ± 2.24 µM compared to 5.87 ± 0.37 µM in AAV-injected WT control mice at P90 (p<0.0001, t-test) (Fig. 3e). Upon brain-wide *Aldh5a1* restoration, AAV-PHP.eB-ALDH5A1-injected Hom mice traveled 1949 ± 135 cm compared to 1808 ± 222 cm as in AAV-FLnP-injected WT control (ns, t-test) (Fig. 3f). These results demonstrate that Hom mice are rescuable using an AAV-mediated gene delivery approach.

**Fig. 3.**
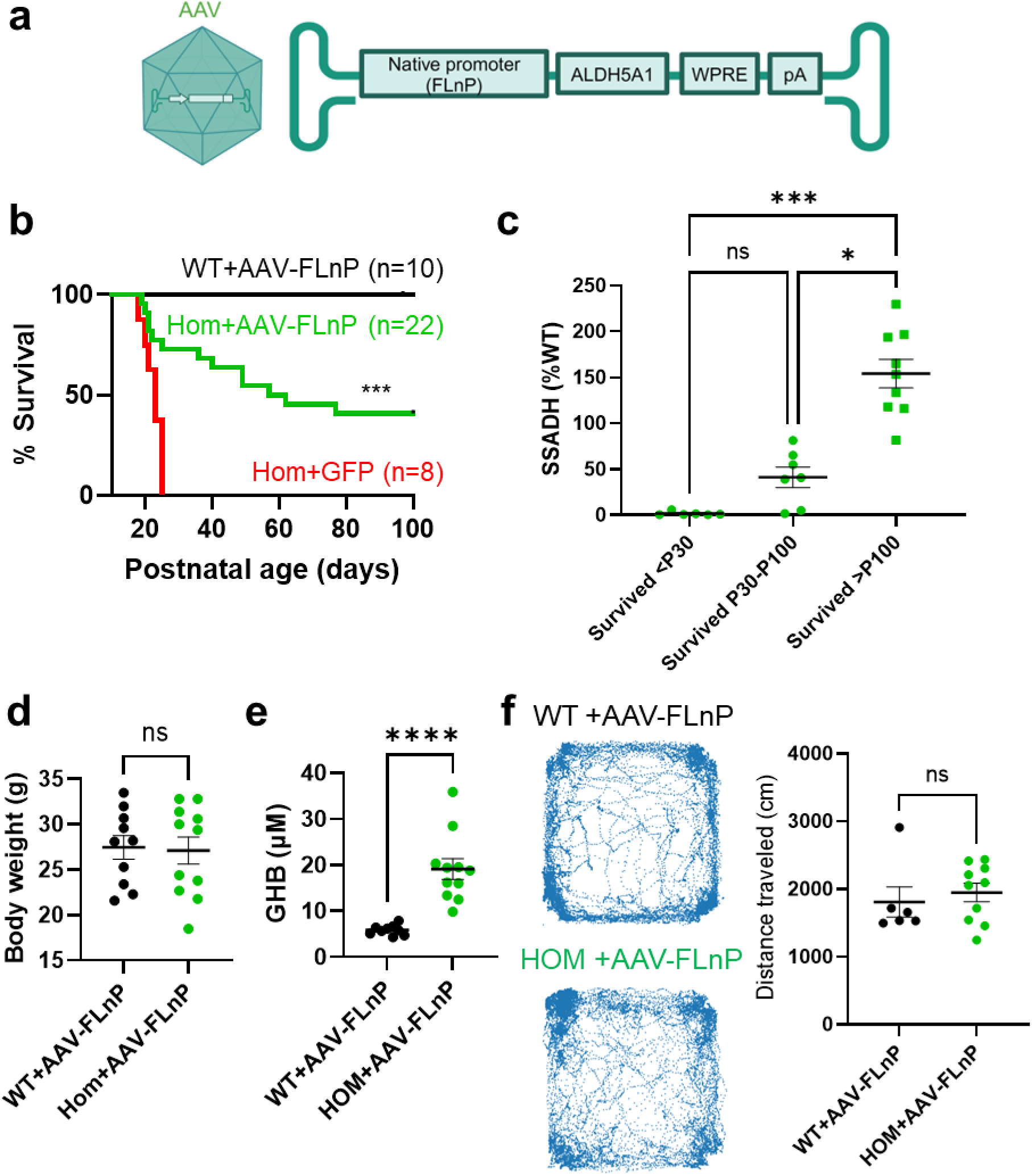
Testing AAV gene therapy candidate for SSADHD in *Aldh5a1^lox-STOP^* mice. (a) Schematic showing the construct payload design of an AAV encompassing a native promoter (FLnP), a full-length ALDH5A1 coding sequence, Woodchuck Hepatitis Virus Posttranscriptional Regulatory Element (WPRE), and a polyadenylation (pA) signal (AAV-FLnP-hALDH5A1, in short, AAV-FLnP). (b) Survival curve of WT or Hom *Aldh5a1^lox-STOP^* mice injected with AAV-FLnP or AAV-GFP control vector, injected at P16. (c) Western blot analysis of Hom *Aldh5a1^lox-STOP^* mice upon AAV-FLnP rescue. The amount of SSADH expressions in their cortex are plotted as a function of their days of survival (<P30, P30-P100, >P100). (d) The body weight of AAV-FLnP-injected WT and Hom at P100 are compared. Individual mouse data points are shown. (e) Serum GHB levels in WT and Hom treated with AAV-FLnP, measured at P100. (f) Free dwelling and exploratory behavior of WT and Hom mice treated with AAV-FLnP, measured at P100. Value = MEAN ± SEM. ****p<0.0001, ns=not significant, log-rank test in (b), One-way ANOVA followed by post-hoc multiple group comparisons in (c). Mann-Whitney test in (d), (e), and (f).

## DISCUSSION

### Preclinical development of a gene replacement therapy for SSADHD, a rare severe neurodevelopmental disorder

In mice modeling complete SSADH loss and its restoration, we address the threshold matter of whether the SSADHD phenotype is reversible with postnatal gene restoration. Encouragingly, we find that either Cre-mediated gene reactivation or AAV-mediated gene replacement reverses essential elements of the model phenotype. Specifically, we identify that brain SSADH protein restoration to >40% WT level corresponds to improved growth and survival, reduced hyperactivity, and reduced blood GHB concentration. The relationship between the magnitude of SSADH restoration and phenotype reversal is underscored by a direct correlation between brain SSADH protein and likelihood of survival. Thus, in this proof-of-concept study, we identified that 1) AAV-mediated brain-wide and peripheral gene restoration approach may be safe and tolerable, and 2) symptom reversal is possible after symptom onset.

SSADHD is a rare disease without effective treatment. Multiple drug trial failures highlight the therapeutic challenge in this complicated condition ^9,15,17,19^. Since no drug therapy is available for SSADHD ^19,28,32–35^, gene replacement may be a realistic option for this patient population ^36^. A liver-directed gene transfer approach in SSADH KO mice conferred limited survival but did not reverse the overall phenotype ^37^. Hepatic SSADH restoration in these studies corresponded to reduced (by as much as 70%) brain and serum GHB concentration ^37^, suggesting an appreciable but incomplete therapeutic potential of peripheral SSADH restoration. The limited success with hepatic *Aldh5a1*gene replacement points to the brain as a major GHB source, and thereby also to brain SSADH restoration as a necessary component of viable SSADHD gene therapy. For this reason, we used the AAV.PHP.eB capsid which enables SSADH expression to be restored simultaneously in the brain and in the periphery. Given recent advances in virus-mediated brain targeting for gene transfer ^38^, eventual clinical SSADHD treatment by simultaneous brain and peripheral organ gene replacement appears realistic.

### Translational path for gene replacement therapy in SSADHD and related GABA disorders

Our results offer a practical insight into a translational path for SSADH-targeted gene therapy. Most importantly, they indicate that SSADH symptoms may be reversible, even if treatment commences at a distinctly symptomatic postnatal age, which would approximate the human scenario where the SSADHD diagnosis is made only after a patient’s symptoms become apparent to the family and treating physicians. In our study, mice were treated at P14-16, which approximates toddler or pre-adolescent human age ^39^. AAV-rescued mice, either after Cre-mediated restoration or treatment with a replacement construct, had a comparable body weight with WT littermates, and did not exhibit an overt hyperactivity phenotype, suggesting that the SSADHD disease trajectory is modifiable by gene replacement therapy even at a symptomatic disease stage.

We also found that blood GHB concentration declines with successful treatment, and correlates directly with symptom relief upon *Aldh5a1* restoration which indicates a role for blood GHB as a mechanistic biomarker and measure of both target engagement and therapeutic efficacy in SSADHD gene therapy clinical trials ^40,41^. Rescued mice had much reduced GHB levels (>98% reduction). Notably, although AAV-rescued mice had GHB levels ∼1.5-fold higher than WT, the residual amount falls within the normal human range of < 49 µM (a recommended cut-off level for GHB levels in normal serum commonly used in forensic studies^42^), suggesting that this heightened GHB level is tolerable.

Last, we achieved a phenotype reversal with systemic delivery of a blood-brain barrier penetrating AAV capsid (i.e., AAV-PHP.eB). While this capsid does not cross the blood-brain barrier in humans, our results indicate that successful SSADHD gene replacement therapy may be achieved by next-generation capsids that enable both systemic and brain gene replacement in humans ^38,43,44^.

### A protein replacement threshold for symptom reversal in SSADHD

A practical question for the translation of SSADH gene replacement therapy is whether a minimal SSADH restoration threshold exists for appreciable therapeutic effects. Since all SSADHD carriers (e.g., patients’ parents) are heterozygous and asymptomatic, 50% of SSADH activity appears tolerable in humans. With AAV-Cre-mediated SSADH restoration to ∼70% WT level, we observed a striking 89% survival of treated mutant mice in contrast to 0% survival in those littermates treated with AAV-GFP control. Western blot analysis from AAV-FLnP-hALDH5A1-treated Hom mouse cortex also revealed a positive correlation between SSADH protein amount and survival: restoration of SSADH above 80% WT resulted in nearly 100% survival beyond P100 (i.e., >5x of untreated Hom mice). This high survival rate has not been achieved by any pharmacotherapy in an SSADH-deficient mouse model previously reported ^21,28,33,35,45^. Interestingly, after treatment by the AAV-FLnP-hALDH5A1 construct, we also observed SSADH levels as high as 200% WT, suggesting a tolerable range of restoration, in addition to the requirement for >50% WT protein as a goal, and an overall broad range of SSADH tolerance upon gene replacement. Beyond the scope of these preliminary experiments, dose-finding experiments with explicit testing of SSADH restoration thresholds, across organs, will help to define a therapeutic range for ALDH5A1 restoration (i.e., replacement SSADH levels that are sufficient to restore GABA and GHB homeostasis, yet are not in excess of tolerable)^22^.

### Therapeutic time window for SSADHD rescue

The developmental trajectory of brain cells and circuits includes critical periods and intrinsic checkpoints beyond which certain cell and circuit functions are no longer modifiable ^46^. In SSADHD, disease-relevant pathologic processes are detectable as early as the embryonic stage ^47^, raising the possibility and concern that prolonged high-GABA and GHB exposure may lead to irreversible brain changes. Fortunately, our data indicates that when the SSADH-encoding gene is delivered at a symptomatic stage, P14, approximating human early childhood, when the mice are markedly underweight, hyperactive, and tremulous, essential aspects of the mouse SSADHD model (i.e., mortality, GHB and gross behavior) can be reversed. Whether other phenotype aspects such as social behavior, learning and memory are also restored to normal will be the work of future studies, beyond the scope of this report.

We anticipate that our SSADHD mouse model will enable future study of phenotype reversibility across ages. Given the critical role of synaptic GABA in early development ^46,48^, whether renormalization of GABA homeostasis (via SSADH restoration) may reverse adverse neuronal network effects due to high-GABA exposure in early life can now be investigated, particularly with Cre-mediated restoration. Furthermore, since the impact of prolonged exposure to GHB ^49^ might be associated with lasting neurological consequences, which can now be studied in detail using our inducible animal model.

### A gene therapy candidate for SSADH deficiency

For clinical translation, we directly tested an AAV construct encompassing a functional native promoter tethered with a functional coding sequence of the *ALDH5A1* gene, delivered systemically using a capsid that has effective blood-brain barrier penetrance. The rationale behind this approach is that SSADH is expressed broadly in the brain and in peripheral organs, particularly liver, and systemic delivery covering brain-wide cellular targets is therefore necessary. In addition, the SSADH expression is cell-specific, so a promoter driving an endogenous expression pattern, constrained to SSADH-expressing cell populations such as inhibitory interneurons and astrocytes in the brain would be ideal. We tested this idea using the native promoter (∼1.6 kb upstream of the human *ALDH5A1* transcriptional start site), coupled with a full length of the human *ALDH5A1* coding sequence. This approach yielded substantial rescue in the SSADH model mice, mimicking the outcomes of *Cre*-dependent SSADH restoration (a controlled, on-demand gene restoration to its endogenous form. Furthermore, we found that the survival of SSADH mutant mice correlates with the amount of SSADH restored. Taken together, these results indicate realistic prospects for SSADHD gene therapy.

### Study limitations

We are cognizant that a range of metrics will require further analysis before a claim of full phenotype reversal is valid. Among these are measures of EEG and seizure activity (since half of the patients with SSADHD have epilepsy), learning and memory and social behavior, as well as circadian and sleep metrics (since patients with SSADHD often have disordered sleep ^50^). At the molecular level, given the altered GABA-A receptor expression profile when GABA and GHB are elevated, future studies can also test whether expression of GABA-A receptor subunits is affected by the disease and restored by therapy. Lastly, our model approximates the most severe SSADHD variant (with uniform early mortality, suggesting that mice do not tolerate SSADH loss as well as humans ^1^), and a milder disease model with some retained enzyme activity and an assessment of its response to gene therapy treatment would add to the preclinical SSADHD field.

## Conclusion

We report the development of an inducible SSADH deficiency mouse model, which allows Cre-tests of dependent SSADH restoration and SSADH restoration by a gene therapy candidate construct. Using this novel mouse model, we demonstrated that the SSADH-associated phenotype is reversable even at a symptomatic stage. For clinical translation, we constructed an AAV vector encompassing a native promoter and coding sequence of the human SSADH encoding gene *ALDH5A1*, packaged into a blood-brain barrier penetrating capsid, delivered systemically. We found that this approach effectively restores SSADH in the mouse brain, with survival correlating to the level of SSADH restoration and GHB normalization, providing a framework for future translation and eventual clinical studies.

## MATERIALS & METHODS

### Study design

We developed a Cre-dependent inducible *Aldh5a1^lox-STOP^* mouse model to study the response of gene replacement therapy in SSADHD. We used AAV-Cre-mediated methods for *Aldh5a1* restoration in *Aldh5a1^lox-STOP^* mice. For phenotypic measures, we included body weight tracking, survival, native behavior, whole blood GHB analysis, western blot analysis, immunohistochemistry, and RNA-seq. A small pilot of male and female comparison was performed but no difference was found between sexes in terms of protein expression, survival, and body weight phenotype. So subsequent experiments were performed in both sexes. All experimental procedures, and materials including mouse lines, viruses, and DNA plasmids are described and approved by IACUC and IBC protocols in Boston Children’s Hospital.

### Construction of Aldh5a1^lox-STOP^ mice

We used the Easi-CRISPR approach to insert a 2k cassette into the first intron of the *Aldh5a1* gene of the mouse. This 2k cassette encompasses a pair of loxP sites, flanking the endogenous splice adaptor immediately upstream of *Aldh5a1* exon two, and a coding sequence for reverse tetracycline transactivator (rtTA) with STOP codons in all frames. A cocktail of 0.61 pmol/μL each of crRNA+tracrRNA, 100 ng/μL Cas9 protein ^51^ and 10 ng/μL ssDNA (IDT) or ds-Donor DNA was injected into the pronuclei of E0.5 embryos (C57Bl6/Hsd). Post-injection survived embryos were re-implanted into recipient CD1 pseudo-pregnant females and allowed to develop to term. The resultant pups were genotyped and genome-edited founders identified and mated to establish lines.

ssDNA megamer used for CRISPR/Cas9 insertion (1999 bp):

ggagctaaccctgaacttctgatcatcctatgctgggatgactgttgtgtgccacaagcccagtttgtgggagctggggattgaacacagggagtcgtacatgcttggcaaactctctaccaaataatccacatccctagtttaagaagaatagacttactctcatctgcctggataaccatgcaatgcaataagcaaacctagcatgtcattctgccattgtaaccacgcagaagtctacttttataattacatttgttaacagagagccagtgggcacacagaacactgttgtctctggagactgtcccttttctaacaagctgcatcttttcacatctcaaaatttttgctgaggataaatcttcaggaacagatagatgccagcactgccttgaaagtaagggcattgctatccttataacttcgtatagcatacattatacgaagttatacgcgtgcagtgaaaaaaat gctttatttgtgaaatttgtgatgctattgctttatttgtaaccattataagctgcaataaacaagttcgttcattagcctggcagcatatcgagatcgaagtcatcgagagcgtcagcagggagcatatccagatcaaaatcatccagggcatcggcgggcagcatgtccaggtcgaagtcgtccagggcgtcggtgggtccgccgctctcgcacttcagctgcttctcgaggccgcagatgatcagttccaggccgaacaggaaggcgggctcggcgccctgtctgtcgaacagctcgatggcctgcttcagcagggggggcatgctgtcggtggtgggtgtctctctttcctctttggcgacctggtgttcctgttcttccagcacgcagcccagggtgaagtggcccacggcgctcagggcgtacagggcgttttccaggctgaagccctgctggcacaggaaggccagctggttttccagtgtctcgtactgcttctcggtgggtctggtgcccaggtgcactttggcgccgtcccggtggctcagcagggcgcatctgtagctcttggcgttgttccgcaggaagtcctgccagctctcgccttccagagggcagctgtgggtgtggtgccggtccagcatctcgatgggcagggcgtccagcagggcccgcttgttcttcacgtgccagtacagggtgggctgttccacgcccagtttctgggccagcttccgggtggtcaggccctcgatgcccacgccgttcagcagttccagggcgctgttgatcactttgctcttgtccagccggctcataggaccggggttttcttccacgtctcctgcttgctttaacagagagaagttcgtggcctggaaaagaaatgaggcagagaaacaatgaactgggaagtgcaggaaagataggttggcacccttgccctcctaagcaccggttacgtgtctagccttacacttggaggaagaatagttcacctcaagtgtcaacttcaccaggctgggggatgccccagtagctagtaaacgttttggattggtttatgggtgttttataacttcgtatagcatacattatacgaagttatacgcgtcgcaggatttcacctattctcatttcataaaacgccaagtgttctccatccagatgagagagaaaacgtattaaaatccaggaggcctggttctgagaaatgactatgactctgtctgtgggcctggagatgtgattcagttggtaaagtgcttccctgacatgcatgaagccctgggtcgcatctttaacacagcataaatctgtgcatggcagaacatgactaatcccagcaccagggagctgtaggcagaacgatcagaagttcaagggactcactgactacataccaagatgccagcttgggctacatagggtcctgtcttagggcaaaagtgaatggcttgctatgtgctgtgaacagggtaccatgcttagtcagagaaagagtgagagaatgtaacatagc

Primer pairs for genotype (checking rtTA-insertion):

rtTA-F: AAGAGCTACAGATGCGCCCT
rtTA-R: GTCTCTCTTTCCTCTTTGGCGA

Primer pairs for genotype (checking Aldh5a1 intron integrity):

Aldh5a1-F: GGACGATTAGTCCAGCCCAG
Aldh5a1-R: CAGGAGGATTGCCTTAGCCC

### Molecular cloning and Sanger sequencing

To confirm the sequence integrity of the Easi-CRISPR inserted construct, the primer pair Aldh5a1-F and Aldh5a1-R were used to capture ∼2kb genomic sequence (Fig. S1a). PCR fragments were then cloned via TOPO cloning, and transformed into E. Coli for clonal selection, followed by Sanger sequencing.

### Molecular cloning of AAV-FLnP-hALDH5A1

The functional native promoter (FLnP) of human *ALDH5A1* sequence (1568 bp) is cloned using the following primers from a DNA construct received from Dr. Patrizia Malaspina at University Tor Vergata, Rome, Italy.

FLnP-F: AAACGCGTACCACTGCACTCCACCC
FLnP-R: AAGAATTCAATCTAGACAACGACGGCGACAGGAAAC

The coding sequence of human *ALDH5A1* (1608 bp) is cloned using the following primers from a DNA construct received from Dr. Gajja Salomons at Department Genetic Metabolic Diseases, University Medical Center at Amsterdam, the Netherlands.

ALDH5A1-F: AATCTAGAGCCGCCATGGCTACATGTATTTGGC
ALDH5A1-R: AAGAATTCCTACAAGCCCCCGTAACACACA

The FLnP-hALDH5A1 payload is subsequently subcloned into the vector pAAV-hSyn-EGFP (Addgene 50465) via restriction enzyme removal of the hSyn promoter and EGFP transgene, to form the AAV-FLnP-hALDH5A1 construct, with its full sequence as follows:

CATGTCCTGCAGGCAGCTGCGCGCTCGCTCGCTCACTGAGGCCGCCCGGGCGTCGGGCGACCTTTGGTCGCCCGGCCTCAGTGAGCGAGCGAGCGCGCAGAGAGGGAGTGGCCAACTCCATCACTAGGGGTTCCTGCGGCCGCACGCGTACCACTGCACTCCACCCTGGGTGACAGAATGAGTCTGTCGGTCTCACAAGAACAAAACAAACAAAAAACAAAAACCACCACCAAAAACAAAACAAAACAAAAACAAAAACTCCCTCCGCAGGCAGGAGATGTAAACACGCCAGCTCCCCAGCCCCACCTGTGATACCTACTGGCCCACATCCCTATTAGGACTCCCTGGTTCTCCCCAGTTCCTGCAATAAAATACAGAGCAAAATAGTGTGAAATTGATTAAACTCTACTGTTATCACTTCTGCCATATGTGACTGGCTCAAAACTCAATGGATCTGACTCAGCAAAAGACAAAGTCAAAATTTAAAGTTACTGCCTCGGCATTTTGCAAATGTCCTAGAAAAAAATTCAACGTCCTAGAAAAAGGTGAAAGCTAGCAAGATACTCTCAACGCTGTAGGAGTTGAGAAAGGGAGATATGTAACCTAAAATCAATGCATAGGGCTGGGCTTCATTTGCACCCACAAATTCCCAGCATACGCCTGCTATTTTTATTTTTATCCTCATTAGGTTTTAAGCTTCCCTGCTGCTTTCAGAGTCTTAAGCCCAAGTCTAATTTTTTAACGGATGAAGAATGCTGTTTAAAGGTGCTCAAATACCTTGGTTGGATAGTATGCTCAATCCTTATACCTTGGTTGATTCAATTTTGACAAATGCTTCCATTTACTAGTTGGTGACCCAACTTTAATCTTACTAGTGGCTTGACTAGGGTAGCTAGTATGACTTTGAACGCTATATTGAATTTAATGTGGCAACTGTAATGAGCATTTTACGTTATCTCACAACAAAATGGGGTAGATATGGTTCTATTTTATAAGTGACTGGACTAAGGTTAACCTGTCGGTCACACAGCTTGTCGGCAACAGACTTGAGTTTGGATCCTAGTCTGTTTATTTCCAAAG CGCTGTTATTTATCGGGAGCAACCCTAGGAGAATGCCTAGACACACACCCAAAAAAGCAGCCAGGCAGCAGAACGCGGGGTCACACTTCGACCCCTCAGAGAACGATCGCTCCCAATCAATACTACGTGCTCAGGAGCTACAACTGAAGAAGCGTGATCACGGCCTTGGCATTTAGGGCAGGCACCAGCGCATCTATGGACCGCGCACAAATTCCGGGGATGCCGAATTTGGGGGATGCTAAGGGGAGAGAGTGGGTCTCTAGCAGCGATTGGGGGCTCAGGAGCAGTTAGTGACAAATGAGCACCCGAAAAGTGAAAAGGTGACAGCAGTCCGCAGGTGCATCTACTGGCGAGCCTTCTCCATCCCCGAACCCAACCCTCCCCCGGGAGAAGGTCGCGCCAGGAGAGAAGCCGCGCGGCGCTTAGGGCAAGGTGCAGAGGGCGGCGCGGCGGTGCAGCGAGAAAGACGCGGAGAGAGGGCGCTGCTCTGTGGCTCTGCAACCTTCCGCCAGCTCCCACGCTTTCCCCGCGCGTCCCCGGCGCCTCCTCGCTCCTCTTGCTTCCCCGCGACCCCTGCGTTCCCGTGCGCGCGCGCCCGCTTGCCTGTTTCCTGTCGCCGTCGTTGTCTAGAGCCGCCATGGCTACATGTATTTGGCTTCGAAGTTGTGGAGCTCGACGACTTGGATCTACATTTCCAGGATGTCGACTTCGACCTCGAGCTGGAGGACTTGTTCCTGCTTCTGGACCTGCTCCTGGACCTGCTCAACTTCGATGTTATGCTGGACGACTTGCTGGACTTTCTGCTGCACTTCTTCGAACAGATAGTTTTGTTGGAGGACGATGGCTTCCTGCTGCTGCAACATTTCCTGTTCAAGATCCTGCTAGTGGAGCTGCTCTTGGAATGGTAGCTGATTGTGGAGTTCGAGAAGCTCGAGCTGCTGTTCGAGCTGCTTATGAAGCATTCTGCCGATGGAGAGAAGTCTCCGCCAAGGAGAGGAGTTCATTACTTCGGAAGTGGTACAATTTAATGATACAAAATAAGGATGACCTTGCCAGAATAATCACAGCTGAAAGTGGAAAGCCACTGAAGGAGGCACATGGAGAAATTCTCTATTCCGCCTTTTTCCTAGAGTGGTTCTCTGAGGAAGCCCGCCGTGTTTACGGAGACATTATCCACACCCCGGCAAAGGACAGGCGGGCCCTGGTCCTCAAGCAGCCCATAGGCGTGGCTGCAGTCATCACCCCGTGGAATTTCCCCAGTGCCATGATCACCCGGAAGGTGGGGGCCGCCCTGGCAGCCG GCTGTACTGTCGTGGTGAAGCCTGCCGAAGACACGCCCTTCTCCGCCCTGGCCCTGGCTGAGCTTGCAAGCCAGGCTGGGATTCCTTCAGGTGTATACAATGTTATTCCCTGTTCTCGAAAGAATGCCAAGGAAGTAGGGGAGGCAATTTGTACTGATCCTCTGGTGTCCAAAATTTCCTTTACTGGTTCAACAACTACAGGAAAGATCCTGTTGCACCACGCAGCAAACTCTGTGAAAAGGGTCTCTATGGAGCTGGGCGGCCTTGCTCCATTTATAGTATTTGACAGTGCCAACGTGGACCAGGCTGTAGCAGGGGCCATGGCATCTAAATTTAGGAACACTGGACAGACTTGTGTTTGCTCAAACCAATTCTTGGTGCAAAGGGGCATCCATGATGCCTTTGTAAAAGCATTCGCCGAGGCCATGAAGAAGAACCTGCGCGTAGGTAATGGATTTGAGGAAGGAACTACTCAGGGCCCATTAATTAATGAAAAAGCGGTAGAAAAGGTGGAGAAACAGGTGAATGATGCCGTTTCTAAAGGTGCCACCGTTGTGACAGGTGGAAAACGACACCAACTTGGAAAAAATTTCTTTGAGCCTACCCTGCTGTGCAATGTCACCCAGGACATGCTGTGCACTCATGAAGAGACTTTCGGGCCTCTGGCACCAGTTATCAAGTTCGATACAGAGGAGGAGGCTATAGCAATCGCTAACGCAGCTGATGTTGGGTTAGCAGGTTATTTTTACTCTCAAGACCCAGCCCAGATCTGGAGAGTGGCAGAGCAGCTGGAAGTGGGCATGGTTGGCGTCAACGAAGGATTAATTTCCTCTGTGGAGTGCCCTTTTGGTGGAGTGAAGCAGTCCGGCCTTGGGCGAGAGGGGTCCAAGTATGGCATTGATGAGTATCTGGAACTCAAGTATGTGTGTTACGGGGGCTTGTAGGAATTCGATATCAAGCTTATCGATAATCAACCTCTGGATTACAAAATTTGTGAAAGATTGACTGGTATTCTTAACTATGTTGCTCCTTTTACGCTATGTGGATACGCTGCTTTAATGCCTTTGTATCATGCTATTGCTTCCCGTATGGCTTTCATTTTCTCCTCCTTGTATAAATCCTGGTTGCTGTCTCTTTATGAGGAGTTGTGGCCCGTTGTCAGGCAACGTGGCGTGGTGTGCACTGTGTTTGCTGACGCAACCCCCACTGGTTGGGGCATTGCCACCACCTGTCAGCTCCTTTCCGGGACTTTCGCTTTCCCCCTCCCTATTGCCACGGCGGAACTCATCGCCGCCTGCCTTGCCCGCTG CTGGACAGGGGCTCGGCTGTTGGGCACTGACAATTCCGTGGTGTTGTCGGGGAAATCATCGTCCTTTCCTTGGCTGCTCGCCTATGTTGCCACCTGGATTCTGCGCGGGACGTCCTTCTGCTACGTCCCTTCGGCCCTCAATCCAGCGGACCTTCCTTCCCGCGGCCTGCTGCCGGCTCTGCGGCCTCTTCCGCGTCTTCGCCTTCGCCCTCAGACGAGTCGGATCTCCCTTTGGGCCGCCTCCCCGCATCGATACCGAGCGCTGCTCGAGAGATCTACGGGTGGCATCCCTGTGACCCCTCCCCAGTGCCTCTCCTGGCCCTGGAAGTTGCCACTCCAGTGCCCACCAGCCTTGTCCTAATAAAATTAAGTTGCATCATTTTGTCTGACTAGGTGTCCTTCTATAATATTATGGGGTGGAGGGGGGTGGTATGGAGCAAGGGGCAAGTTGGGAAGACAACCTGTAGGGCCTGCGGGGTCTATTGGGAACCAAGCTGGAGTGCAGTGGCACAATCTTGGCTCACTGCAATCTCCGCCTCCTGGGTTCAAGCGATTCTCCTGCCTCAGCCTCCCGAGTTGTTGGGATTCCAGGCATGCATGACCAGGCTCAGCTAATTTTTGTTTTTTTGGTAGAGACGGGGTTTCACCATATTGGCCAGGCTGGTCTCCAACTCCTAATCTCAGGTGATCTACCCACCTTGGCCTCCCAAATTGCTGGGATTACAGGCGTGAACCACTGCTCCCTTCCCTGTCCTTCTGATTTTGTAGGTAACCACGTGCGGACCGAGCGGCCGCAGGAACCCCTAGTGATGGAGTTGGCCACTCCCTCTCTGCGCGCTCGCTCGCTCACTGAGGCCGGGCGACCAAAGGTCGCCCGACGCCCGGGCTTTGCCCGGGCGGCCTCAGTGAGCGAGCGAGCGCGCAGCTGCCTGCAGGGGCGCCTGATGCGGTATTTTCTCCTTACGCATCTGTGCGGTATTTCACACCGCATACGTCAAAGCAACCATAGTACGCGCCCTGTAGCGGCGCATTAAGCGCGGCGGGTGTGGTGGTTACGCGCAGCGTGACCGCTACACTTGCCAGCGCCTTAGCGCCCGCTCCTTTCGCTTTCTTCCCTTCCTTTCTCGCCACGTTCGCCGGCTTTCCCCGTCAAGCTCTAAATCGGGGGCTCCCTTTAGGGTTCCGATTTAGTGCTTTACGGCACCTCGACCCCAAAAAACTTGATTTGGGTGATGGTTCACGTAGTGGGCCATCGCCCTGATAGACGGTTTTTCGCCCTTTGACGTTGGAGTCCACGTTCTTTAATAGTGGACTCT TGTTCCAAACTGGAACAACACTCAACTCTATCTCGGGCTATTCTTTTGATTTATAAGGGATTTTGCCGATTTCGGTCTATTGGTTAAAAAATGAGCTGATTTAACAAAAATTTAACGCGAATTTTAACAAAATATTAACGTTTACAATTTTATGGTGCACTCTCAGTACAATCTGCTCTGATGCCGCATAGTTAAGCCAGCCCCGACACCCGCCAACACCCGCTGACGCGCCCTGACGGGCTTGTCTGCTCCCGGCATCCGCTTACAGACAAGCTGTGACCGTCTCCGGGAGCTGCATGTGTCAGAGGTTTTCACCGTCATCACCGAAACGCGCGAGACGAAAGGGCCTCGTGATACGCCTATTTTTATAGGTTAATGTCATGATAATAATGGTTTCTTAGACGTCAGGTGGCACTTTTCGGGGAAATGTGCGCGGAACCCCTATTTGTTTATTTTTCTAAATACATTCAAATATGTATCCGCTCATGAGACAATAACCCTGATAAATGCTTCAATAATATTGAAAAAGGAAGAGTATGAGTATTCAACATTTCCGTGTCGCCCTTATTCCCTTTTTTGCGGCATTTTGCCTTCCTGTTTTTGCTCACCCAGAAACGCTGGTGAAAGTAAAAGATGCTGAAGATCAGTTGGGTGCACGAGTGGGTTACATCGAACTGGATCTCAACAGCGGTAAGATCCTTGAGAGTTTTCGCCCCGAAGAACGTTTTCCAATGATGAGCACTTTTAAAGTTCTGCTATGTGGCGCGGTATTATCCCGTATTGACGCCGGGCAAGAGCAACTCGGTCGCCGCATACACTATTCTCAGAATGACTTGGTTGAGTACTCACCAGTCACAGAAAAGCATCTTACGGATGGCATGACAGTAAGAGAATTATGCAGTGCTGCCATAACCATGAGTGATAACACTGCGGCCAACTTACTTCTGACAACGATCGGAGGACCGAAGGAGCTAACCGCTTTTTTGCACAACATGGGGGATCATGTAACTCGCCTTGATCGTTGGGAACCGGAGCTGAATGAAGCCATACCAAACGACGAGCGTGACACCACGATGCCTGTAGCAATGGCAACAACGTTGCGCAAACTATTAACTGGCGAACTACTTACTCTAGCTTCCCGGCAACAATTAATAGACTGGATGGAGGCGGATAAAGTTGCAGGACCACTTCTGCGCTCGGCCCTTCCGGCTGGCTGGTTTATTGCTGATAAATCTGGAGCCGGTGAGCGTGGGTCTCGCGGTATCATTGCAGCACTGGGGCCAGATGGTAAGCCCTCCCGTATCGTAGT TATCTACACGACGGGGAGTCAGGCAACTATGGATGAACGAAATAGACAGATCGCTGAGATAGGTGCCTCACTGATTAAGCATTGGTAACTGTCAGACCAAGTTTACTCATATATACTTTAGATTGATTTAAAACTTCATTTTTAATTTAAAAGGATCTAGGTGAAGATCCTTTTTGATAATCTCATGACCAAAATCCCTTAACGTGAGTTTTCGTTCCACTGAGCGTCAGACCCCGTAGAAAAGATCAAAGGATCTTCTTGAGATCCTTTTTTTCTGCGCGTAATCTGCTGCTTGCAAACAAAAAAACCACCGCTACCAGCGGTGGTTTGTTTGCCGGATCAAGAGCTACCAACTCTTTTTCCGAAGGTAACTGGCTTCAGCAGAGCGCAGATACCAAATACTGTTCTTCTAGTGTAGCCGTAGTTAGGCCACCACTTCAAGAACTCTGTAGCACCGCCTACATACCTCGCTCTGCTAATCCTGTTACCAGTGGCTGCTGCCAGTGGCGATAAGTCGTGTCTTACCGGGTTGGACTCAAGACGATAGTTACCGGATAAGGCGCAGCGGTCGGGCTGAACGGGGGGTTCGTGCACACAGCCCAGCTTGGAGCGAACGACCTACACCGAACTGAGATACCTACAGCGTGAGCTATGAGAAAGCGCCACGCTTCCCGAAGGGAGAAAGGCGGACAGGTATCCGGTAAGCGGCAGGGTCGGAACAGGAGAGCGCACGAGGGAGCTTCCAGGGGGAAACGCCTGGTATCTTTATAGTCCTGTCGGGTTTCGCCACCTCTGACTTGAGCGTCGATTTTTGTGATGCTCGTCAGGGGGGCGGAGCCTATGGAAAAACGCCAGCAACGCGGCCTTTTTACGGTTCCTGGCCTTTTGCTGGCCTTTTGCTCA

### AAV packaging and administration

Cre-mediated cassette removal in *Aldh5a1^lox-STOP^* mice was conducted by IP injection at P16 using pre-packaged pAAV-CAG-Cre-WPRE-hGH developed at BCH as described previously ^52^. The pre-packaged virus was available from the Viral Core at BCH.

### Genetic mouse model

TdTomato (Ai14) mice were obtained from JAX (#007914), which is a characterized Cre-dependent reporter tool strain harboring a loxP-flanked STOP cassette preventing transcription of a red fluorescent protein variant (tdTomato) driven by a CAG promoter ^53^.

### Western blot

Protein expression was determined using western blotting as described ^22^. Brain tissues (cortex) were lysed by a buffer containing 0.22% Beta glycerophosphate, 0.18% Sodium orthovanadate, 5% Sodium deoxycholate, 0.38% EGTA, 1% Sodium lauryl sulfate, 6.1% Tris, 0.29% EDTA, 8.8% Sodium chloride, 1.12% Sodium pyrophosphate decahydrate, 10% ethoxylated Nonylphenol, and protease inhibitors, with debris removed by centrifugation. Samples were resolved and transferred onto a nitrocellulose membrane. After blocking, protein blots were incubated with antibodies against SSADH. Anti-β-actin was used as a loading control. Chemiluminescence signals were detected by LI-COR imaging and quantified using the Image Studio software. All data was collected with the experimenter blinded to sample experimental conditions.

| Item name | Source / Catalog # |
| --- | --- |
| AAV-PHP.eB-CAG-Cre-WPRE-hGH | BCH Viral Core |
| AAV-PHP.eB-FLnP-hALDH5A1 | BCH Viral Core, UMass Med School |
| Anti-SSADH | Abcam / ab129017 |
| Anti-β actin | Abcam / ab6276 |

### Cryosectioning and confocal imaging

Under deep anesthesia, mice were perfused transcardially with ice-cold PBS followed by 4% paraformaldehyde (PFA). Brain tissues were harvested and postfixed in 4% PFA for 24 h at 4°C before transferring into 30% sucrose. Cryopreserved brain tissue was then frozen in Tissue-Plus OCT Compound (Fisher Healthcare) and stored at -80°C for > 24 h before sectioning. Free-floating cryosections (coronal, 30 µm) were obtained at -20°C, washed briefly with PBS, and incubated with primary antibodies overnight at 4°C followed by Alexa Fluor-conjugated secondary antibodies for 1 h at RT. Slides were sealed with mounting medium with DAPI (DAPI Fluoromount G, Southern Biotech) and glass coverslips and stored at 4°C until imaging with the LSM700 microscope (Zeiss). All data was collected with the experimenter blinded to sample experimental conditions.

### Whole blood GHB analysis

GHB quantitation was performed as previously described ^54^. Whole blood from freshly dissected mice was spotted on filter paper. GHB extraction was performed using 200 µL of methanol containing 500 ng/mL d6GHB internal standard added on a blood-spotted paper. Aqueous extract was injected into an HPLC-MS/MS system in partial loop injection with needle overfill mode. Ions were formed by electro-spray ionization in negative ion mode and detected with multiple reaction monitoring (MRM). Blood GHB content was quantified by Targetlynx software. A calibration curve was determined using GHB/d6GHB. All data was collected with the experimenter blinded to sample experimental conditions.

### Native exploratory behavior

A mouse was placed inside in containment chamber containing an optically transparent footpad sensor, and a high-speed bottom-up facing camera ^30^. The footpad sensor used frustrated total internal reflection (FTIR) technology, employing near-infrared light to selectively illuminate the inferior body surfaces in contact with the sensor, such that the contact area of the paws varies according to the relative weight borne by individual paws and the forces exerted by individual toes and footpads within each paw, to provide a real-time readout of foot position, pressure, form, and gait. Separate data streams representing changes in footprints and body form were differentially analyzed to reveal bodily movement and behaviors. Each chamber is 20cm x 20cm x 20cm and made of black acrylic for easy sanitization between recording sessions. One mouse was evaluated in a chamber at one time. All data were collected with experimenter blinded to sample experimental conditions.

### Statistics

Mann Whitney nonparametric test and one-way ANOVA was used for two-group and multiple-group comparisons, respectively, in general. Group survival curve was assessed using log-rank test. Specific tests are all indicated in corresponding figure legends. Group sample size was derived from preliminary or prior studies, based on variance of available data sets and a default power value of 0.8. Statistical significance was designated as p<0.05.

## Supporting information

SSADH mice before treatment

SSADH mice after AAV-Cre

## Availability of data and materials

All data, code, and materials used in the analysis are available in main text, supplementary materials, or raw data format to any researcher from the corresponding author upon reasonable request for purposes of reproducing or extending the analysis. The use of *Aldh5a1^lox-STOP^* mice might be subject to materials transfer agreement (MTA) under Boston Children’s Hospital policy.

## Acknowledgement

We thank the IDDRC Gene Manipulation Core, funded by NIH P50 HD105351.

## Author contributions

HHCL, AR conceptualized this study. HHCL, CJW, GG, AR provided methodology. HHCL, GM, AL, ZZ, BW, SAV, EA, RL, MN, DC performed investigation. HHCL provided visualization of data. HHCL, PLP, MS, AR acquired funding. HHCL, AR administered the project. HHCL, TY, PLP, CJW, GG, MS, AR supervised the project. HHCL, AR contributed to the original draft of this manuscript. HHCL, GM, AL, ZZ, BW, SAV, EA, RL, MN, DC, TY, CJW, PLP, GG, MS, AR reviewed and edited this manuscript.

## DECLARATIONS

## Funding

National Institutes of Health grant 1R21NS121858 (HHCL, AR), 1R61NS131704 (HHCL, AR, MS)

SSADH Association Research Program (HHCL, AR, PLP, MS)

Boston Children’s Hospital Translational Research Program (HHCL, AR)

## List of Supplementary Materials

Materials and Methods

Fig. S1 to 4

Movie S1 – *Aldh5a1^lox-STOP^* mice and control littermate before treatment (at P20)

Movie S2 – *Aldh5a1^lox-STOP^* mice and control littermate after AAV-Cre treatment (at P100)

References (1–54)

---

**Fig. S1.**
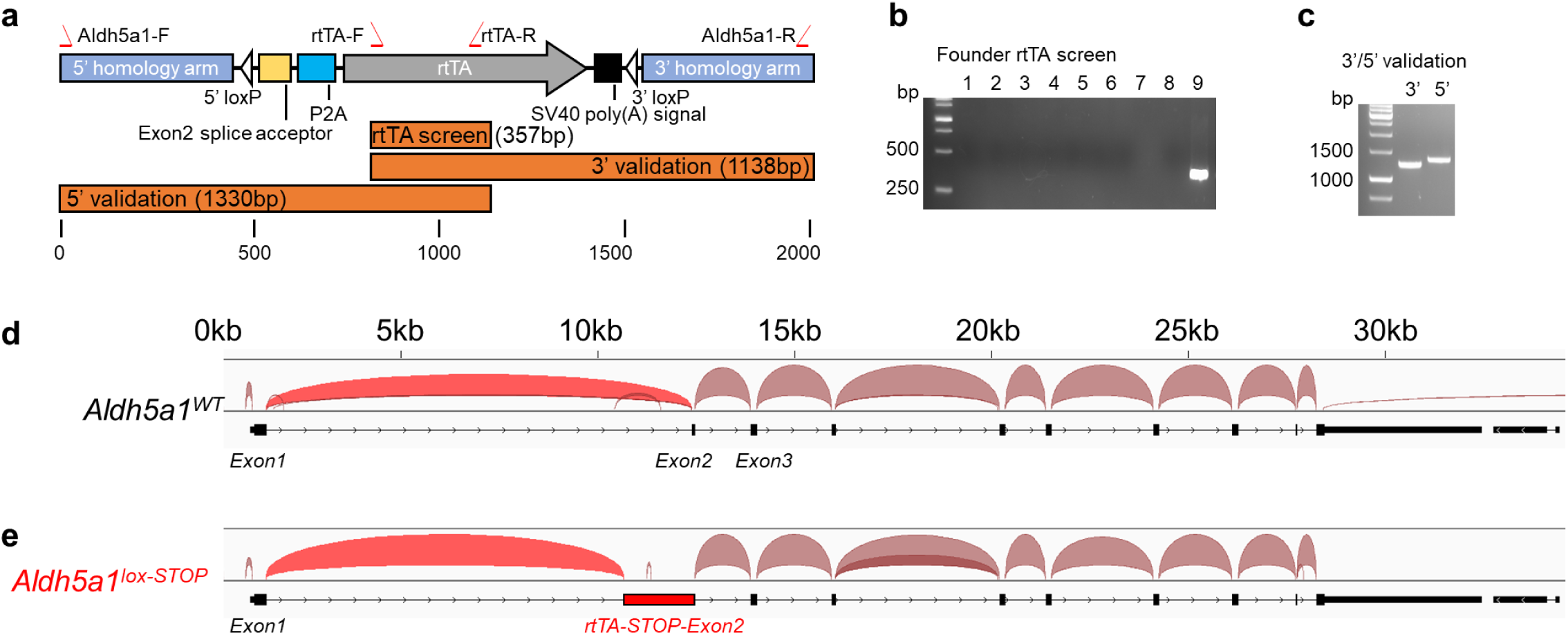
Molecular design of Aldh5a1^lox-STOP^ mice, genotyping verification, and RNA-seq validation of obligated slicing of the endogenous *Aldh5a1* gene. (a) The inserted Lox-STOP-Lox cassette, encompassing an exon two splice acceptor site, a P2A, a reverse tetracycline transactivator (rtTA), an SV40 poly(A) signal, and multiple STOP codons covering all frames. Locations of primers are indicated, for various PCR genotyping conditions. (b) A DNA gel showing a positive band of ∼357bp in animal #9 upon PCR of the rtTA fragment. DNA ladder reference is shown on the left. (c) A DNA gel showing the PCR validation experiments using two sets of independent primers for 3’ and 5’ amplifications of the Lox-STOP-Lox cassette. (d-e) Mapping analysis of RNA-seq results demonstrating the readings across exons in WT allele and lox-STOP allele (instead of exon two, the transcripts are mapped to the rtTA-STOP exon2.

**Fig. S2.**
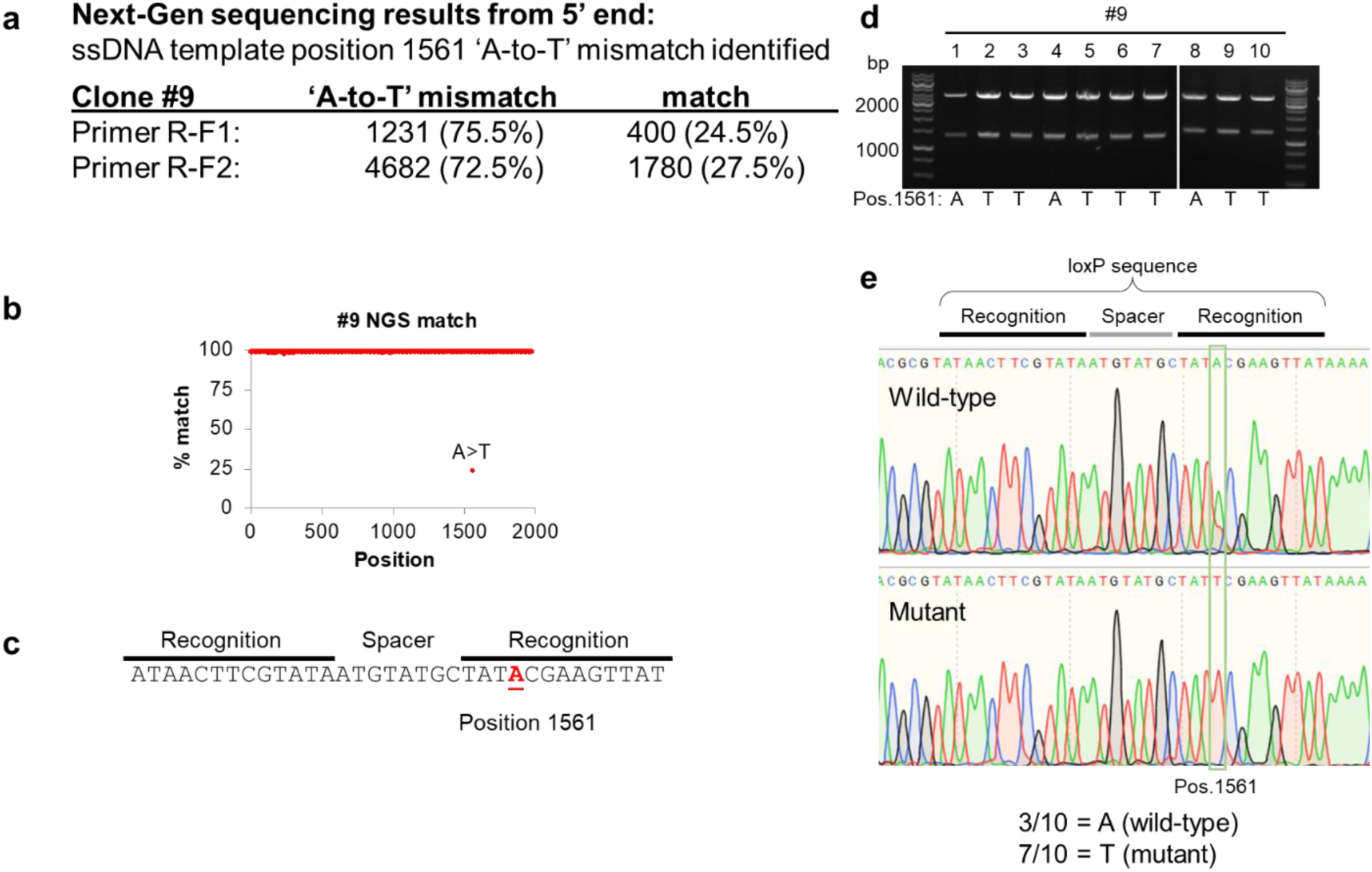
Next-Gen sequencing revealed one single nucleotide replacement in the inserted sequence of Aldh5a1^lox-STOP^ mice. (a) One nucleotide ‘A-to-T’ mismatch at position 1561 of the inserted sequence was identified. (b) the mismatch position is shown. (c) the mismatch position is found within the recognition sequence of loxP site. (d-e) Sanger sequencing confirming the mismatch.

**Fig. S3.**
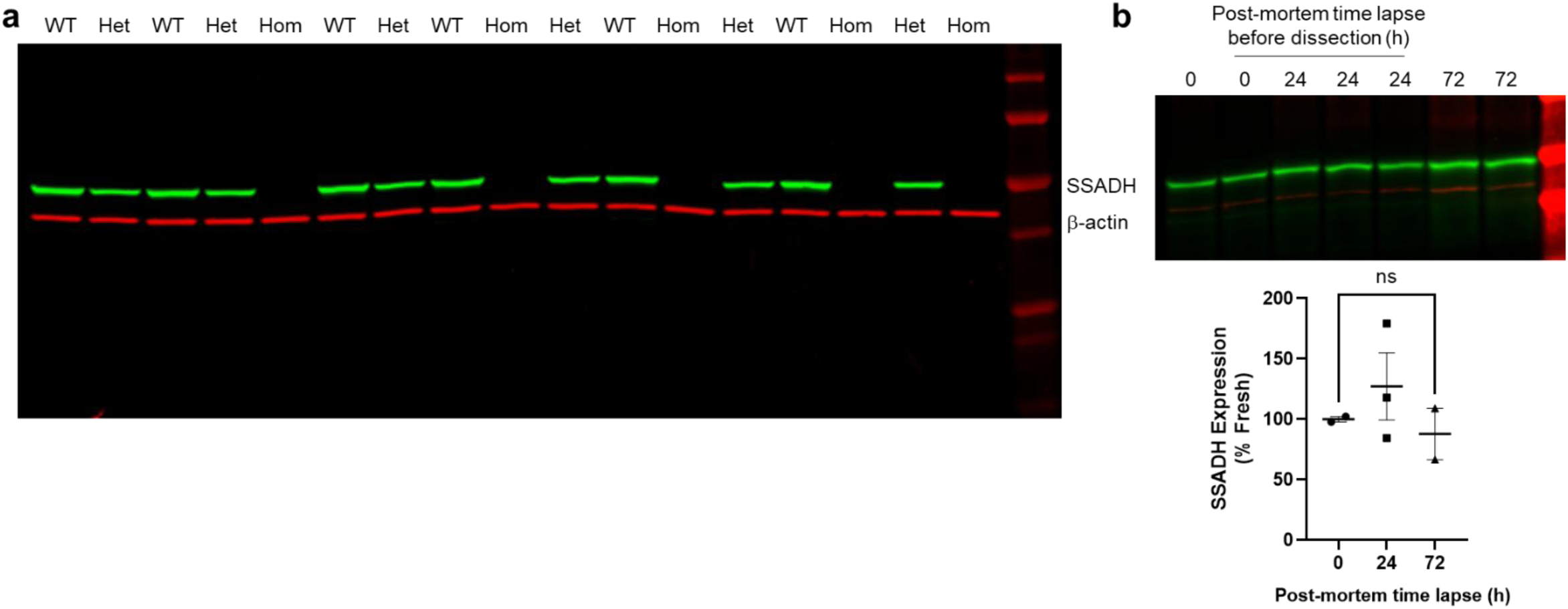
SSADH protein is not subjected to post-mortem degradation. (a) Western blot showing the SSADH (in green) and β-actin (in red) of cortical lysates obtained from WT, Heterozygous (Het), and Homozygous (Hom) *Aldh5a1^lox-STOP^* mice at P16-18. (b) Western blot analysis shows the expression of SSADH proteins in the cortex at 0, 24, and 72 hours (h) collected post-mortem. The β-actin is used as the loading control. Mean and SEM are shown. ns= not significant, One-way ANOVA followed by Tukey’s multiple comparisons test.

**Fig. S4.**
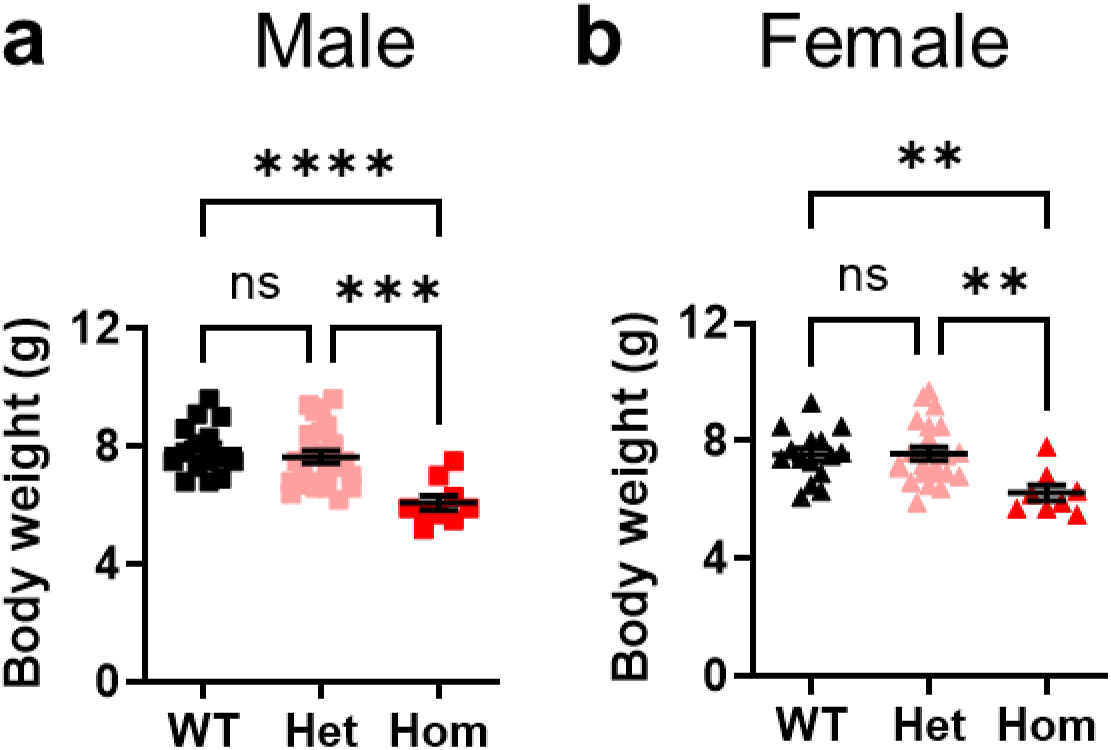
Body weight of Aldh5a1^lox-STOP^ mice. (a) Individual male mouse body weight of WT, Het, and Hom *Aldh5a1^lox-STOP^* mice at P16. (b) Individual female mouse body weight of WT, Het, and Hom *Aldh5a1^lox-STOP^* mice at P16. Mean and SEM are shown. **p<0.01, ****p<0.0001, ns = not significant, One-way ANOVA followed by post-hoc multiple group comparisons.

